# Identification and characterization of domesticated bacterial transposases

**DOI:** 10.1101/119735

**Authors:** Frederic Bertels, Jenna Gallie, Paul B Rainey

## Abstract

Selfish genetic elements (SGEs), such as insertion sequences (ISs) and transposons are found in most genomes. Transposons are usually identifiable by their high copy number within genomes. In contrast REP associated tyrosine transposases (RAYTs), a recently described class of bacterial transposase, are typically present at just one copy *per* genome. This suggests that RAYTs no longer copy themselves and thus they no longer function as a typical transposase. Motivated by this possibility we interrogated thousands of fully sequenced bacterial genomes in order to determine patterns of RAYT diversity, their distribution across chromosomes and accessory elements, and rate of duplication. RAYTs encompass exceptional diversity and are divisible into at least five distinct groups. They possess features more similar to housekeeping genes than insertion sequences, are predominantly vertically transmitted and have persisted through evolutionary time to the point where they are now found in 24% of all species for which at least one fully sequenced genome is available. Overall, the genomic distribution of RAYTs suggests that they have been co-opted by host genomes to perform a function that benefits the host cell.

## Introduction

Mechanisms for the maintenance of selfish genetic elements (SGEs) have been discussed for over 40 years (Nevers & Saedler 1977; Doolittle & Sapienza 1980; Charlesworth et al. 1994). To persist in the long term, the subsequent host generation must contain more SGE copies on average than the previous host generation (Burt & Trivers 2006). This can be achieved even when the element confers a fitness cost, either through mechanisms that disproportionately increase copy number through self-replication and horizontal gene transfer (Doolittle & Sapienza 1980), or by mechanisms that kill offspring devoid of the SGE (*e.g.*, meiotic drive or toxin-antitoxin (TA) systems (Gerdes et al. 2005; Naito et al. 1995; Hurst et al. 1996)).

While SGEs need not benefit the host (Hickey 1982; Bichsel et al. 2013), many SGEs carry genes that do enhance host fitness. In some cases SGEs have even been domesticated. That is, they have evolved to perform functions that provide direct benefit to the host organism. Examples of such fitness enhancing functions include adaptive immunity, where genes derived from transposases mediate rearrangement of gene cassettes involved in antigen recognition (Jones & Gellert 2004), stress response systems derived from bacterial TA systems (Van Melderen & De Bast 2009; Van Melderen 2010), and defence systems against foreign DNA, such as caspases from CRISPRs (Krupovic et al. 2014).

Recently a new class of bacterial transposases has been described: REP associated tyrosine transposases (RAYTs). RAYTs are found in a wide range of bacteria (Nunvar et al. 2010; Bertels & Rainey 2011b; Ton-Hoang et al. 2012), but unlike typical transposases, which are part of insertion sequences (ISs; (Mahillon & Chandler 1998), RAYTs occur as single copy elements. They are characteristically associated with short repetitive extragenic palindromic sequences (REPs) that are typically arranged as pairs of REP sequences termed REPINs (REP doublets forming hairpINs; (Bertels & Rainey 2011a; 2011b). REPINs are non-autonomous mobile elements that are significantly over-represented in many bacterial genomes, and are likely dependent upon RAYTs for their dissemination (Nunvar et al. 2010; Bertels & Rainey 2011b; Ton-Hoang et al. 2012). Hence RAYTs possess the functionality of a transposase but seem unable to replicate. Instead they appear to disperse REPINs throughout bacterial genomes.

Given that RAYTs appear to be present at just a single copy per genome, their maintenance is difficult to explain. If RAYTs cannot copy themselves then they cannot be maintained by selection on transposition activity. This suggests that RAYTs are maintained by virtue of a functional relationship with the host cell, or with REPINs, or via a combination of both (Bertels & Rainey 2011b). It has been argued that because REPs perform distinct functions within bacterial cells, ranging from transcriptional termination (Espéli et al. 2001) to regulation of translation (Liang et al. 2015), RAYTs may be domesticated transposable elements (Ton-Hoang et al. 2012; Siguier et al. 2014). While evidence supports the notion that REP sequences have been co-opted to perform diverse functions, there is little evidence to suggest this is also true for RAYTs. To date the only function assigned to RAYTs is movement of REPINs, but evidence remains indirect (Nunvar et al. 2010; Bertels & Rainey 2011b; Ton-Hoang et al. 2012; Messing et al. 2012).

One possible explanation is that RAYTs are maintained by hitchhiking with beneficial mutations caused by movement of REPINs. This requires that the rate of mutation caused by REPIN movement is high and in the same of order of magnitude as that caused by a defect in the methyl-directed mismatch repair system responsible for mutator genotypes (Sniegowski et al. 2000), but no such evidence exists. In fact recent work calculates that REPINs duplicate approximately every 60 million generations (Bertels et al. 2017). This exceedingly low rate of duplication casts doubt on the possibility that RAYTs are maintained by hitchhiking, thus leaving open the possibility that RAYTs are domesticated transposons (Bertels & Rainey 2011a; 2011b).

Progress toward understanding the causes of RAYT maintenance might be derived from a detailed analysis of the evolutionary characteristics of RAYTs. We report such an analysis here focussing on the pattern of RAYT diversity across thousands of bacterial genomes. Central to our analysis is a comparison of the RAYT family with two families of well-characterized housekeeping genes (*def* and *tpiA*) and two known IS families (IS*200* and IS*110*). We show that RAYTs share characteristics with housekeeping genes rather than with ISs. We also reveal five divergent RAYT types. Our data indicate that RAYTs have no capacity for selfish replication and so, in addition to within-genome dissemination of REPINs, RAYTs likely perform some currently unrecognized function that is central to their maintenance across a broad range of bacterial genomes.

## MATERIAL AND METHODS

### Acquisition of genome sequences

Bacterial genome sequences were downloaded from the NCBI ftp site on the 11^th^ of June 2013 (ftp://ftp.ncbi.nih.gov/genomes/archive/old_genbank/Bacteria/). On that day 2,950 bacterial chromosomes and 2,106 plasmids were fully sequenced and available for analysis.

### Identification of family members by BLAST

For each gene family (RAYTs, IS*200*, IS*100*, *def*, *tpiA*) one arbitrarily chosen protein was used as a query for a BLAST search. The query sequence for the RAYT family was YafM from *Pseudomonas fluorescens* SBW25 (PFLU_RS20900). Query sequences for housekeeping genes were *P. fluorescens* SBW25 peptide deformylase, Def (PFLU_RS00090), and *P. fluorescens* SBW25 triosephosphate isomerase TpiA (PFLU_RS25840). Query sequences for the ISs were ECIAI1_4438 from *Escherichia coli* IAI1 (IS*200* family) and, ISEc*32* from *E. coli* S88 plasmid pECOS88 (IS*110* family, ECSMS35_RS25240). The sequence of each protein was used to interrogate 2,950 chromosomes and 2,106 plasmids using TBLASTN (protein search against nucleotide database).

Search results were analysed as follows. First, all database matches with an e-value of less than 1e-2, 1e-5, 1e-15, and 1e-20 were recorded. Second, for each of the recorded database matches all coding sequences (CDS) from Genbank annotations that overlapped with the match were determined. Third, for overlaps with a single gene, the DNA sequence of the gene and its encoded amino acid sequence were extracted; for multiple overlaps the longest overlapping open reading frame was used for sequence extraction; matches without overlapping genes were ignored. Fourth, in addition to recording the DNA and amino acid sequence of the overlapping gene, the DNA sequence of the 5’ and 3’ extragenic space immediately flanking the identified gene was stored.

### Identification of duplication events (analysis 2)

For all homologs that occur in the same genome, nucleotide sequences were aligned using the Needleman-Wunsch algorithm (Needleman & Wunsch 1970) and pairwise identities (the number of sites identical between two sequences divided by the total number of sites) calculated. We used the standard BLAST nucleotide substitution matrix NUC.4.4. Gap opening cost was set to 6 and gap extension was set to 1. All pairs with an identity greater than 95% were deemed duplicates.

### Determination of genome-wide frequencies of flanking 16-mers (analysis 4)

Frequencies for all oligonucleotides of length 16 (16mers) from all replicons (chromosomes and plasmids) were determined, according to Bertels and Rainey (2011b). In brief, extragenic 5’ and 3’ 16mer frequencies were determined for each family member using a sliding window with a step size of one for both leading and lagging strands. With knowledge of this frequency, the most abundant 16mer from each flanking non-coding DNA sequence could be determined.

### Calculating pairwise identity of amino acid sequences

Pairwise alignments between protein sequences were computed by applying the Needleman-Wunsch algorithm (Needleman & Wunsch 1970). We used BLOSUM65 as substitution matrix. Gap opening cost was set to 10 and gap extension was set to 1. Pairwise identity is the number of identical sites within the alignment divided by the total number of sites.

### Definition of RAYT protein sequence groups

To determine sequence groups a Markov clustering algorithm (MCL) with default parameters was applied to a matrix of pairwise protein identities greater than 26% (Van Dongen 2000). For proteins longer than 80 amino acids an identity threshold of 24.8% has previously been used for identifying homologs (Sander & Schneider 1991). The resulting sequence clusters were visualized with Cytoscape (www.cytoscape.org) (Cline, et al. 2007).

### Phylogenetic analyses

To build a combined RAYT/IS*200* phylogeny, three protein sequences from each RAYT and IS*200* cluster containing more than 30 sequences (six RAYT and six IS*200* groups) were randomly selected. These 36 sequences were then aligned with default parameters on MUSCLE (Edgar 2004) and a phylogeny was generated using PhyML with the BLOSUM62 scoring matrix (Guindon and Gascuel 2003). IS*200* Group 6 was excluded because sequences from this group could not be aligned. We visualized trees with Geneious v10.0.8 (Kearse et al. 2012).

### Phylogenetic congruence analysis at the sub-genus taxonomic level

We determined phylogenetic congruence at the sub genus taxonomic level by comparing the phylogenies of the two concatenated housekeeping genes (*tpiA* and *def*) with each of the five RAYT groups. We chose the RAYT subsets from those bacterial genera where RAYTs are most common. Hence we identified genera that are well represented in the genome database that also contain a large number of RAYTs. For Group 1 we chose *Pseudomonas*, for Group 2 we chose *Escherichia*, for Group 3 *Pseudomonas*, for Group 4 *Haemophilus* and for Group 5 we selected *Pseudomonas*. Nucleotide sequences were aligned with clustalo (Sievers et al. 2011) and reconstructed trees with PhyML using a GTR substitution model (Guindon & Gascuel 2003) for both the concatenated housekeeping genes and the RAYT Groups. For comparison we also reconstructed a phylogeny from IS*110* genes that occur in *Pseuodomonas.* We visualized trees with Geneious v10.0.8 (Kearse et al. 2012).

### Phylogenetic congruence analysis for RAYTs from different species

For the three largest RAYT groups we chose a single RAYT member from all species belonging to the Gammaproteobacteria and that also contain a member of either IS*110* or IS*200* for comparison. These sequences were aligned with clustalo (Sievers et al. 2011), and phylogenetic trees generated by PhyML (Guindon & Gascuel 2003). Comparisons among trees were made using treedist from the phylip suite (Felsenstein 2009).

## RESULTS AND DISCUSSION

### Overview of strategy for comparing RAYTs, ISs and housekeeping gene families

ISs persist by replicating within and between genomes (Doolittle & Sapienza 1980; Bichsel et al. 2013), with dissemination between genomes being mediated by elements capable of horizontal transmission (*e.g.,* plasmids). Hence, ISs are expected to be present in multiple copies within the genomes of individual bacteria, and to be over-represented on transmissible elements such as plasmids. In contrast, genes necessary for bacterial persistence (*e.g.*, housekeeping genes) typically occur in a single copy *per* genome, and are not dependent on horizontal transfer.

In order to assess whether RAYTs show patterns of diversity typical of ISs or housekeeping genes, we first identified members of five gene families and then compared familial characteristics. The five gene families are RAYTs, two IS families (IS*200* and IS*110*), an essential housekeeping gene family (*def*; peptide deformylase), and a non-essential but highly conserved gene family (*tpiA*; triosephosphate isomerase). For each family, we investigated the following four characteristics: (1) the average number of family members *per* replicon (chromosome or plasmid), (2) the proportion of family members for which identical (or nearly identical) copies are present on the same replicon, (3) the proportion of family members present on plasmids, and (4) the average replicon-wide frequency of the most common 16bp sequence from the DNA sequences flanking each family member. The results for the RAYT family were compared to those obtained for the other families.

### Identification of members of the RAYT, IS*200*, IS*110*, *def* and *tpiA* gene families

We defined gene families as groups of genes that share a similar sequence with a single representative of that gene family (reference gene).

IS and housekeeping gene families are of comparable size (Figure 1, **Supplementary Figure 1**). The *def* gene family is the largest (3,841 members; members found in 92% of all bacterial genomes searched and 89% of all species) followed by IS*200* (3,232; members present in 26% of all bacterial genomes searched and 30% of all species), IS*110* (2,806; members found in 26% of all bacterial genomes searched and 29% of all species) and *tpiA* (2,782; members present in 93% of all bacterial genomes searched and 91% of all species). The RAYT family is considerably smaller; containing 1,045 members with members occurring in 24% of all bacterial species for which at least one fully sequenced genome was available.

**Figure 1.**
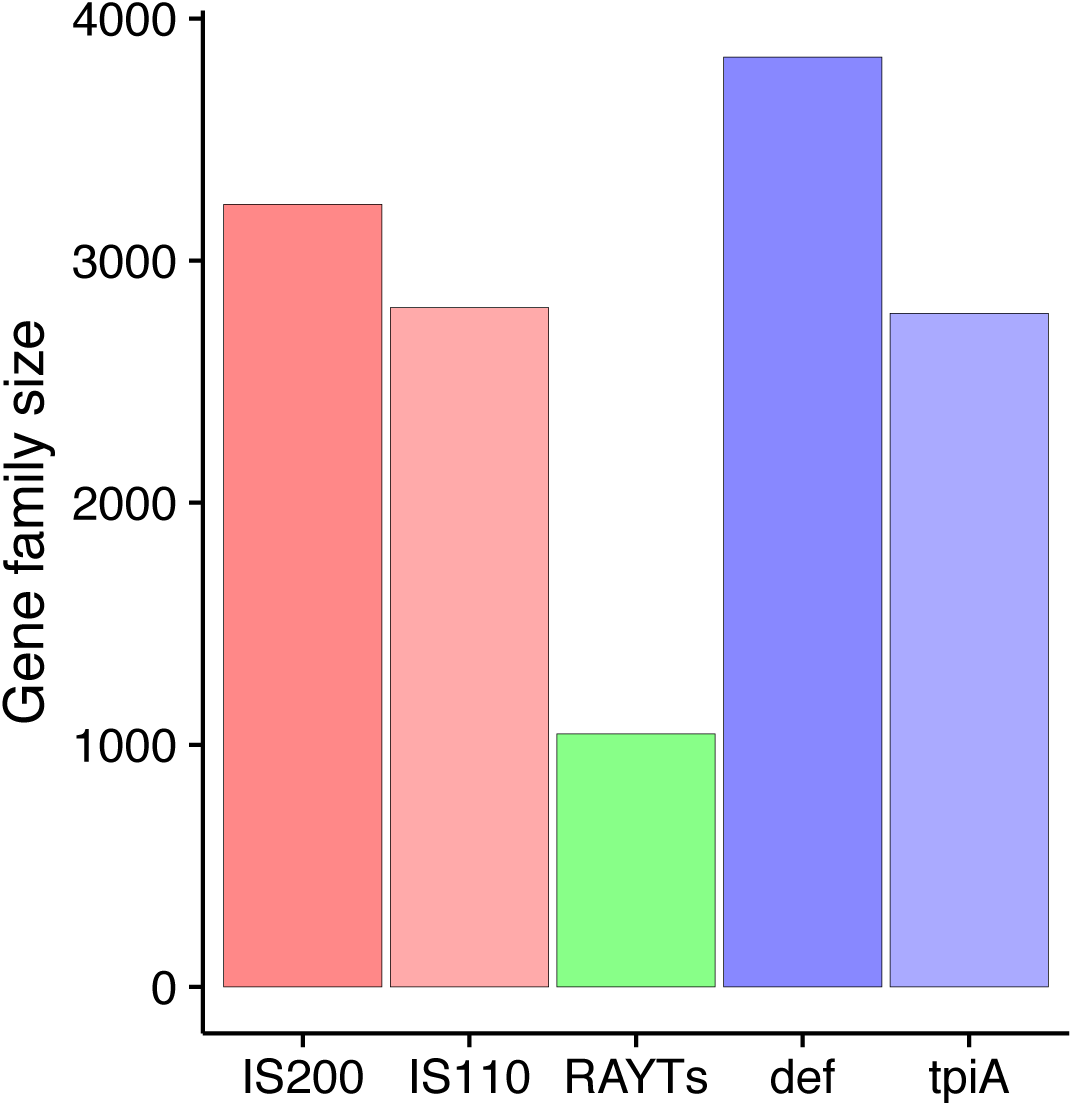
Family size of ISs, RAYTs and housekeeping genes.

### Analysis 1: Number of family members *per* replicon is high for IS elements and low for housekeeping genes

Having determined the number of genes in each family, we next asked how gene family members are distributed across replicons. That is, for replicons containing at least one member of a particular gene family, how many members are present on average (Figure 2A)?

**Figure 2.**
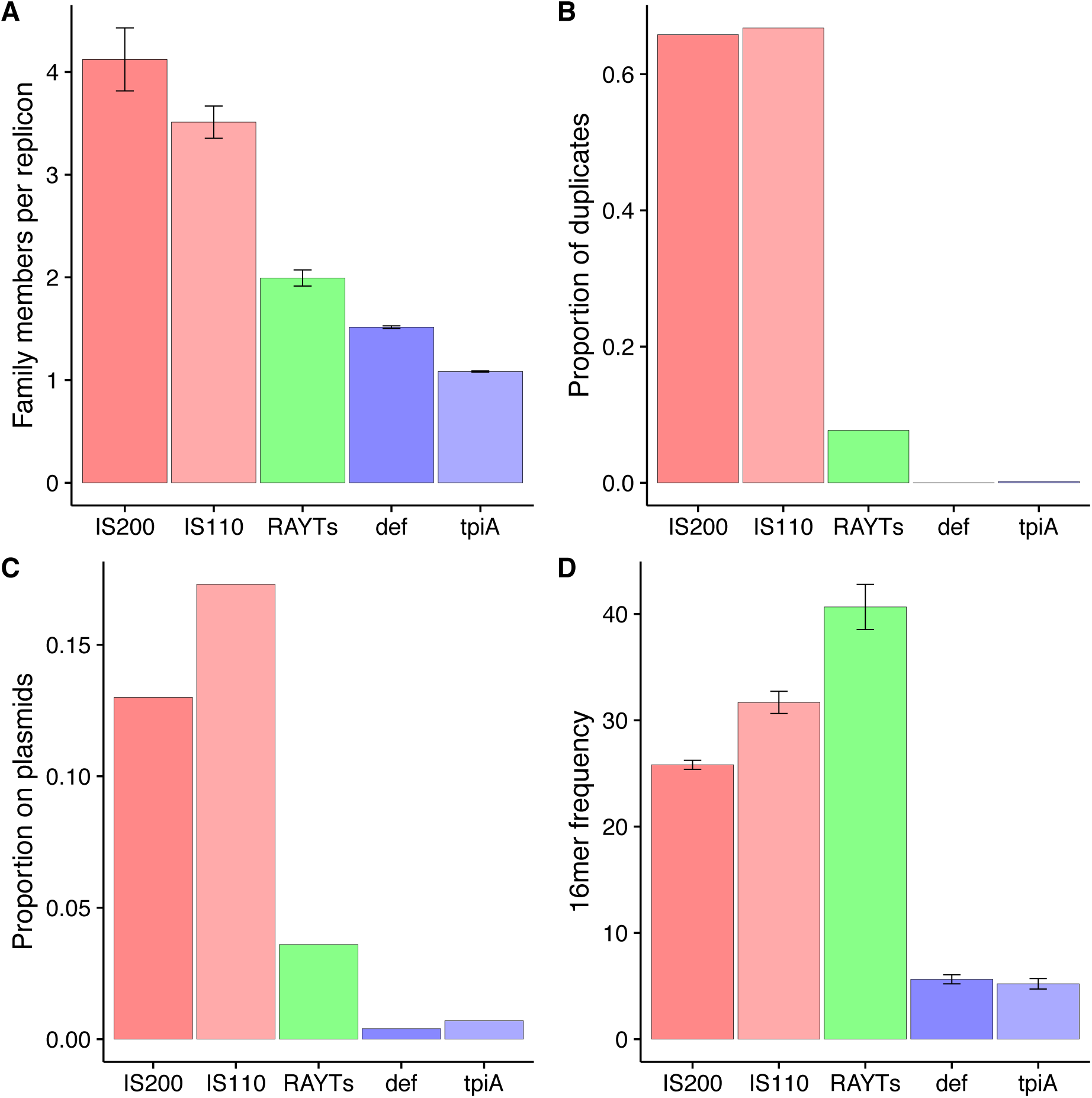
RAYT characteristics are more similar to those of housekeeping genes than IS elements. (*A*) Average copy number of each gene family *per* replicon (excluding replicons in which no family members occur). (*B*) Proportion of genes for which at least one duplicate (pairwise nucleotide identity of ≥95%) was found on the same replicon. (*C*) Proportion of family members found on plasmids. (*D*) Average frequency of the most abundant 16mer identified in the flanking 5′ and 3′ extragenic regions of each family member. All error bars show standard errors.

Replicons that harbour IS*200* or IS*110* contain on average between three and four copies *per* replicon, respectively. For the housekeeping gene family *tpiA* the number of copies *per* replicon is much lower (1.08). For the *def* family the number is slightly higher (1.5). This difference in size between the two housekeeping gene families may have been caused by an ancient duplication event that led to the addition of a family of *def* paralogs (genes derived from ancient duplication events) (Bergthorsson et al. 2007; Näsvall et al. 2012). RAYTs are present at almost two *per* replicon. This hints at either some capacity for replication, or diversification of the RAYT family. We analyse this in detail in later sections.

### Analysis 2: Duplication rates distinguish housekeeping genes from IS elements

Analysis 1 encompasses **all** members of a family present on a replicon. This includes recently duplicated genes, genes derived from ancient duplications (paralogs) and laterally acquired genes. Of these, recently duplicated genes are of greatest interest, because they are likely to have arisen from recent self-duplication events (*i.e.* SGE activity). Accordingly, one would expect the relative size of this gene subset to be considerably larger for SGEs than for genes with a beneficial function. Thus, taking advantage of the fact that recent duplication events will have resulted in the presence of gene copies with identical (or nearly so) DNA sequences, we next determined the proportion of each gene family for which at least one nearly identical copy (≥95% nucleotide sequence identity) exists on the same replicon (Figure 2B).

About 66% of all IS*200* (2127/3232) and IS*110* (1874/2806) genes appear to have been recently duplicated (Figure 2B). For these families, the proportion of duplicated genes correlates with the number of copies *per* replicon (**Supplementary Figure 2A**; F-statistic for a linear fit of data for both IS families *P*=6.5 × 10^-5^). We observe that, with increasing proportion of duplications, there is an increase in the number of gene copies *per* replicon in both IS families. No such correlation exists for either RAYTs or housekeeping genes, as expected from the low proportion of gene duplications (see below). This means that for ISs the change in the average number of family members *per* replicon (**Supplementary Figure 2A**) is not due to ancient duplication events leading to diversification of the IS family; rather, it is due to differences in duplication rates for IS family members more distantly related to the reference gene.

In contrast to ISs, the two housekeeping gene families show almost no evidence of recent duplication. There were no duplications among *def* gene family members (0/3841), and only 0.2% (3/2782) for the *tpiA* family. Of all the RAYT genes, 7.7% (80/1045) were found to have recently duplicated with the additional copy being present on the same replicon. The proportion of RAYT duplication events is less than that observed for IS*200* and IS*110*, but nevertheless higher than that of housekeeping gene families. As we show in later sections, this pattern results from the inclusion of divergent IS*200* sequences in the RAYT family by the BLAST search.

### Analysis 3: Presence on plasmids as a proxy for horizontal transfer rates

In the above analyses, both chromosomes and plasmids were interrogated for the presence of homologous genes. Here we determined the proportion of family members that are found solely on plasmids (Figure 2C). Under the assumption that plasmids facilitate the horizontal transfer of genes, we expect SGEs such as ISs to be present more frequently than housekeeping genes on plasmids (Bichsel et al. 2013).

ISs are commonly found on plasmids; 13% of IS*200* genes and 17.3% of IS*110* genes are found on plasmids. Contrastingly, members of the two housekeeping gene families are almost never found on plasmids - only 0.4% of all *def* genes and 0.7% of all *tpiA* genes occur on plasmids. For RAYTs, 3.6% of family members occur on plasmids. This value is between that obtained for ISs and housekeeping genes. However at the slightly more stringent e-value of 1e-5 in our BLAST search, only 0.3% of all RAYTs occur on plasmids, a value very similar to that found for the housekeeping gene families (**Supplementary Figure 2**). Again as we show below this observation can be explained by the inadvertent inclusion of divergent IS*200* members into the RAYT family at higher e-values.

### Analysis 4: High 16mer frequencies in the flanking extragenic space is a core feature of RAYT genes

Many bacterial genomes contain short, palindromic sequences that are overrepresented in extragenic space. These sequences have been named repetitive extragenic palindromic sequences (REPs) (Higgins et al. 1982; Stern et al. 1984), and have been shown to replicate as part of a doublet termed a REPIN (Bertels & Rainey 2011b). REPINs are almost always found in the extragenic space flanking RAYT genes. These flanking REPINs usually contain highly abundant 16bp long sequences (16mers) (Bertels & Rainey 2011b). We generalize this finding and use the abundance of the most common 16mer of the 5′ and 3′ extragenic flanking regions as a feature to distinguish RAYTs from other genes.

For the IS*200* and IS*110* families, we found higher frequencies of flanking 16mers than for housekeeping genes: the average frequency of flanking 16mers ranged between 25 and 32. The most common flanking 16mer for ISs are typically functionally linked to the IS (Mahillon & Chandler 1998) as they are part of the flanking terminal repeats. Hence, a significant correlation is expected between the proportion of gene family members that are duplicates and 16mer frequencies. The higher the proportion of duplicates the higher the flanking 16mer frequency. Such a correlation is evident from **Supplementary Figures 2D** and **2B** (F-statistic for a linear fit *P*=1.7 × 10^-5^) supporting the notion that the most common flanking 16mer is part of the IS.

We found that 16mers in the adjacent extragenic space of housekeeping genes are repeated infrequently in the remainder of the replicon: average frequencies range from 5.2 to 5.6. For 100 randomly assembled genomes of ∼6.7Mb in length (the length of the *P. fluorescens* SBW25 genome), the most abundant 16mer occurs on average 5.08 times (Bertels & Rainey 2011b). The frequency for 16mers flanking housekeeping genes is only slightly higher than these values, indicating that duplications do not affect flanking 16mer frequencies for housekeeping genes.

The RAYT gene family was found to have the highest frequencies of flanking 16mers across replicons (41). As expected there is no correlation between 16mer frequencies and the proportion of duplicated genes (F-statistic for a linear fit *P*=0.24, ranging from 2% for the most stringent family definition to 7.7%, **Supplementary Figure 2B** and **D**). Thus gene duplication events are not the cause for the observed high frequencies of 16mers flanking RAYT genes. Instead high flanking 16mer frequencies are caused by RAYTs disseminating neighbouring REPINs throughout the genome.

### RAYT protein sequences cluster into distinct groups

In the above analyses, we determined mean gene family characteristics. In the following section we investigate at a finer scale characteristics of RAYTs. To determine whether the RAYT family consists of distinct groups we applied the MCL clustering algorithm (Van Dongen 2000) to a protein similarity matrix of RAYTs. RAYT sequences cluster into at least six groups, each containing 30 or more members (Figure 3A). The groups are ordered by size, i.e. Group 1 contains the most members Group 6 contains the least (sequences are available as **Supplementary Data**).

**Figure 3.**
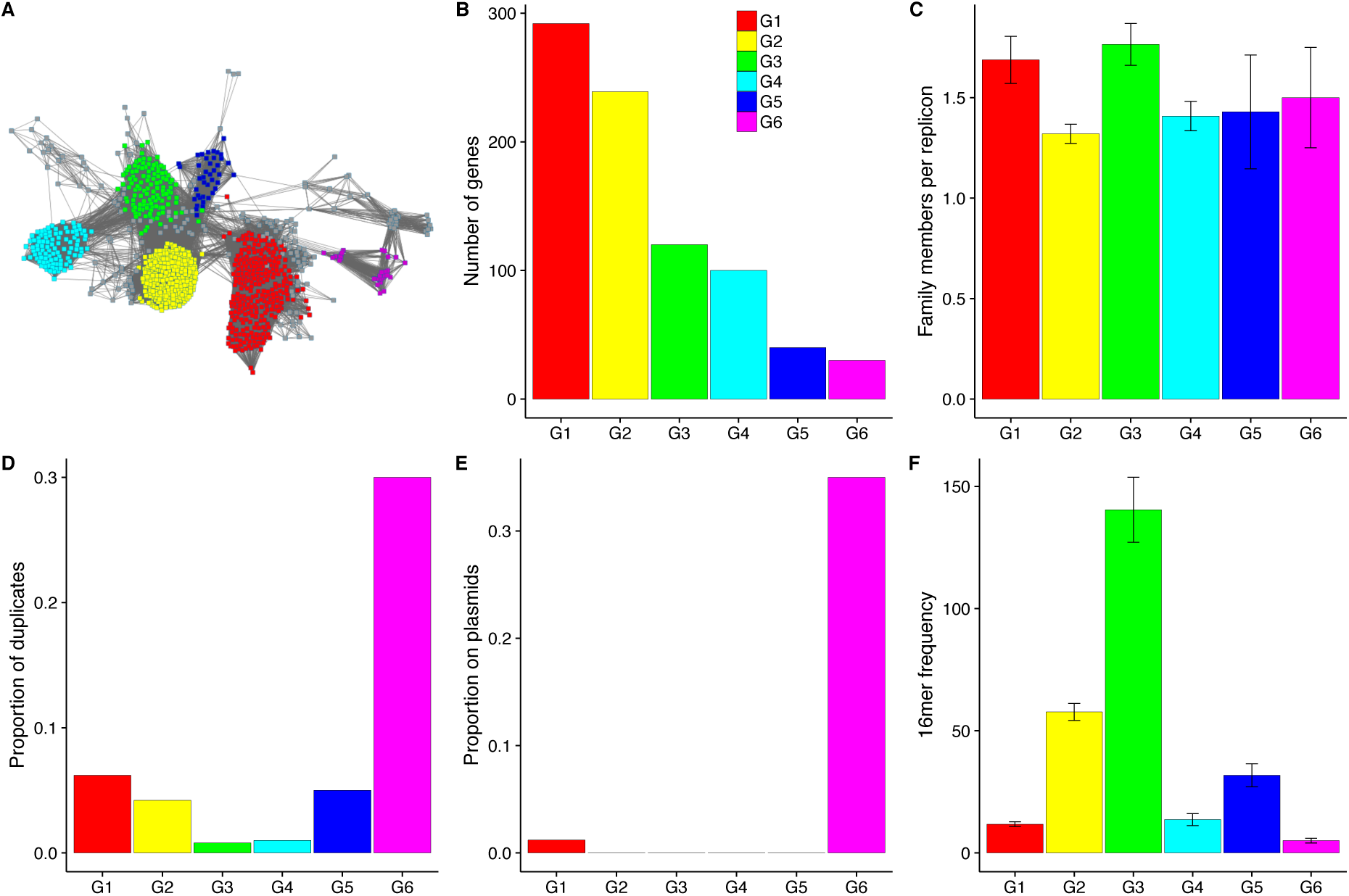
Protein similarity network and genomic characteristics show significant divergence of RAYT groups. (A) RAYT protein relationship map constructed by Cytoscape using the organic layout (Cline et al. 2007). RAYT groups identified by MCL are highlighted according to the legend. Nodes represent RAYT proteins. Nodes are connected if the pairwise amino acid identity is ≥26%. Differences between the Cytoscape protein clustering (proximity of nodes) and the MCL algorithm (colour of nodes) are due to the difference in the information provided to MCL and Cytoscape: Cytoscape only receives information on whether or not nodes are connected, whereas MCL performs coloration based on the exact pairwise similarity between proteins. Graphs (B) to (F) contain the same information as Figures 1 and **2**, but for the six largest RAYT groups in (A) as opposed to the five gene families.

Group 3 contains members of the previously identified Clade I (Bertels & Rainey 2011b). Members of RAYT Clade II defined by Bertels and Rainey, 2011, are found in Group 2. Group 2 also contains *E. coli* RAYTs, for which experimental evidence on REP-RAYT interactions have been determined *in vitro* (Ton-Hoang et al. 2012; Messing et al. 2012). An alignment shows that 12 of the 18 nucleotide binding amino-acid sites are conserved in RAYT Group 2 (**Supplementary Figure 5**) but only one site is conserved across all IS*200* and RAYT groups (**Supplementary Figure 6**), highlighting the large diversity of RAYTs (Bertels et al. 2017; Bertels & Rainey 2011b). Groups 1, 4 and 5 are newly identified RAYT groups.

The mapping of RAYT groups onto host taxonomy is shown in **Supplementary Figure 7**. Groups show various patterns of distribution but no strong overall association with host taxonomy. For example, Group 1 is widely distributed across the Proteobacteria (although rare in the Alphaproteobacteria), but is also found in the Clostridia. Group 4 is rare among the Proteobacteria, but common in the Cyanobacteria, Chlorobia and Caldilineae. Group 1 shows the widest distribution with representatives found across 21 different bacterial classes. However, Group 1 is also by far the largest RAYT group occurring in 150 different bacterial species Although Group 4 occurs in only 61 species it is found in 17 different bacterial classes.

### RAYT protein groups show characteristics similar to housekeeping genes with one exception

For each of the six groups we determined the genomic distribution characteristics as done previously for the five gene families above (Figure 3B-F, **Supplementary Figure 3**). Little variation was observed in the number of group members *per* replicon: the numbers range from 1.3 (Group 2) to 1.8 (Group 3) copies *per* replicon (Figure 3C). Despite this, the proportion of group members with a nearly identical copy on the same replicon differs dramatically between groups (Figure 3D). Group 6 (30%) and Group 1 members (6.3%) have the highest proportion of duplicates, whereas Group 3 has the lowest (0.8%). Surprisingly, although Group 3 has the lowest proportion of duplicates, it is the group with the highest number of gene copies *per* replicon, possibly indicating that this group is more functionally diverse than the other RAYT groups. The fact that there are multiple distinct RAYT genes per genome has been noted previously but has not been quantified (Bertels & Rainey 2011b; Nunvar et al. 2010; Loper et al. 2012).

The proportion of group members found on plasmids shows a similar pattern to the proportion of duplicates observed for each group (Figure 3E). Again, Group 6 shows by far the highest proportion on plasmids (35%). The only other RAYT group for which any members were found on plasmids was Group 1 (1.2%), which also has the second highest proportion of duplicates (Figure 3D). Hence, of all RAYT groups, Group 1 and 6 are the two that most closely resemble ISs in terms of their distribution characteristics.

The maximum frequency of 16mers found in extragenic space indicates which of the RAYT groups is associated with REP sequences (Figure 3F). Only Group 2 (57 replicon-wide occurrences) and Group 3 (140 occurrences) members are associated with 16mers that are more frequently found across replicons than those associated with ISs. However, Group 1 (12 occurrences) and Group 4 (14 occurrences) are associated with 16mers that are more common than 16mers flanking housekeeping genes. Group 5 (31 occurrences), despite not showing signs of high duplication rates, is associated with 16mers that are as frequent as those associated with insertion sequences. Group 6 (5), despite relatively high duplication rates is associated with 16mers that show low abundance in the genome.

The divergence of Group 6 elements from the RAYT family raises the possibility that Group 6 RAYTs might belong to a distinct family of ISs. IS*200* is the most likely close relative (Bertels & Rainey 2011b; Nunvar et al. 2010; Ton-Hoang et al. 2012). To investigate this possibility, the IS*200* family was subjected to identical analyses as the RAYT family.

### IS*200* groups have IS-like characteristics with one exception

The structure of the IS*200* family is much more homogenous than that of the RAYT family (Figures 4A and 3A). All groups are arranged around the largest groups: Group 1 and Group 2. The only subgroup that is distinctly separated is Group 3, a distinction that is also evident in the very different characteristics of Group 3.

**Figure 4.**
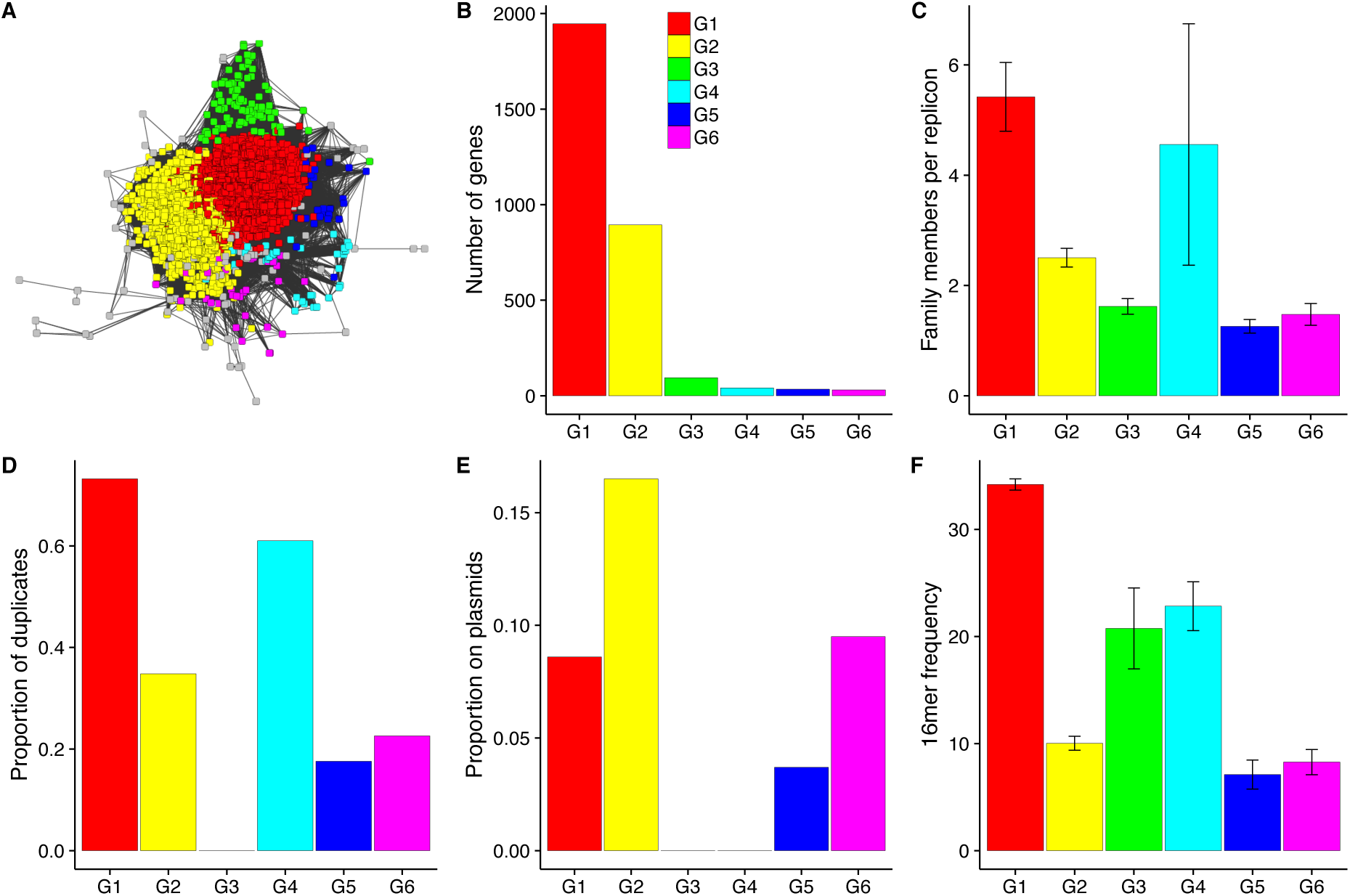
Protein similarity network and genomic distribution of IS*200* groups. (A) shows a protein similarity network of the IS*200* family. The map was produced as described in the legend for Figure 3A. Graphs (B) to (F) show the same genomic characteristics as in Figures 2 and 3.

Group sizes in IS*200* are less evenly distributed than RAYT groups (Figure 4B). Groups 1 and 2 (the central groups in Figure 4A) have by far the most members (1946 and 894), whereas the remaining groups all have less than 100 members.

As expected for ISs, the number of gene copies *per* replicon (Figure 4C) correlates with the observed proportion of duplicates for each group (Figure 4D). The proportion of duplicates is high for all IS*200* groups except for Group 3, for which we did not find a single duplicate. Groups with a high proportion of duplicates have a high number of gene copies *per* replicon (F-statistic for a linear fit for all groups *P=*10^-12^). Again the outlier is Group 3, which has more gene copies *per* replicon than Group 5 and Group 6, but no duplicates. The proportion of genes on plasmids is high for four out of the six IS groups (Figure 4E). As expected from the above results, we did not observe any Group 3 members on plasmids. More surprisingly, we did not observe Group 4 members on plasmids. This is unexpected for an IS with an extremely high replication rate (proportion of duplicates is 61%). To ensure this finding is not an artefact of how the reference was chosen the analysis was repeated using a member of Group 4 as the reference gene. In this analysis Group 4 became part of the main IS*200* cluster. Hence, as also evident in the gene cluster in Figure 4A, Group 4 is not a distinct group but belongs to the main IS*200* gene family. Furthermore, this indicates that the reason the proportion of genes on plasmids is low is an artefact of sampling, as all members of Group 4 in our original analysis occurred on only nine different replicons.

Typical for ISs, the frequency of flanking 16mers again correlates with the proportion of duplicates (**Supplementary Figure 4F**; F-statistic for a linear fit for all groups *P=*10^-6^), except for Group 3, which is associated with highly abundant 16mers (21 occurrences across replicons at 1e-2) but shows no signs of duplication.

Our analyses show that RAYTs and IS*200* sequences have very different characteristics. RAYTs show features more typical of housekeeping genes and IS*200* shows features typical of ISs. However, in each family we identified one group that did not fit into the overall pattern. It is possible that the IS-like RAYT group belongs to the IS*200* family, given its distant relationship to the RAYT family. Similarly, although not as distantly related to the IS*200* query sequence, it is possible that IS*200* Group 3 belongs to the RAYT family. To investigate these possibilities we performed a combined phylogenetic and network analysis for the RAYT and IS*200* Groups.

### The IS-like RAYT subgroup is part of the IS*200* gene family

The combined RAYT and IS*200* phylogeny shows the IS*200* and RAYT families are distinct with the exception of RAYT Group 6, which clusters within the IS*200* phylogeny (Figure 5A). In contrast, the RAYT-like IS*200* Group 3 clusters as a distinct group with the IS*200* family. However, according to the phylogenetic tree, IS*200* Group 3 is the most closely related IS*200* group to the RAYT family. This raises the possibility that RAYTs have been domesticated only once and that the IS*200* Group 3 has maintained sequence features similar to IS*200* sequences. This hypothesis, however, needs to be treated with caution as the high divergence between the different genes, potential convergent evolution and horizontal gene transfer hinders ability to infer the phylogenetic history with confidence.

**Figure 5.**
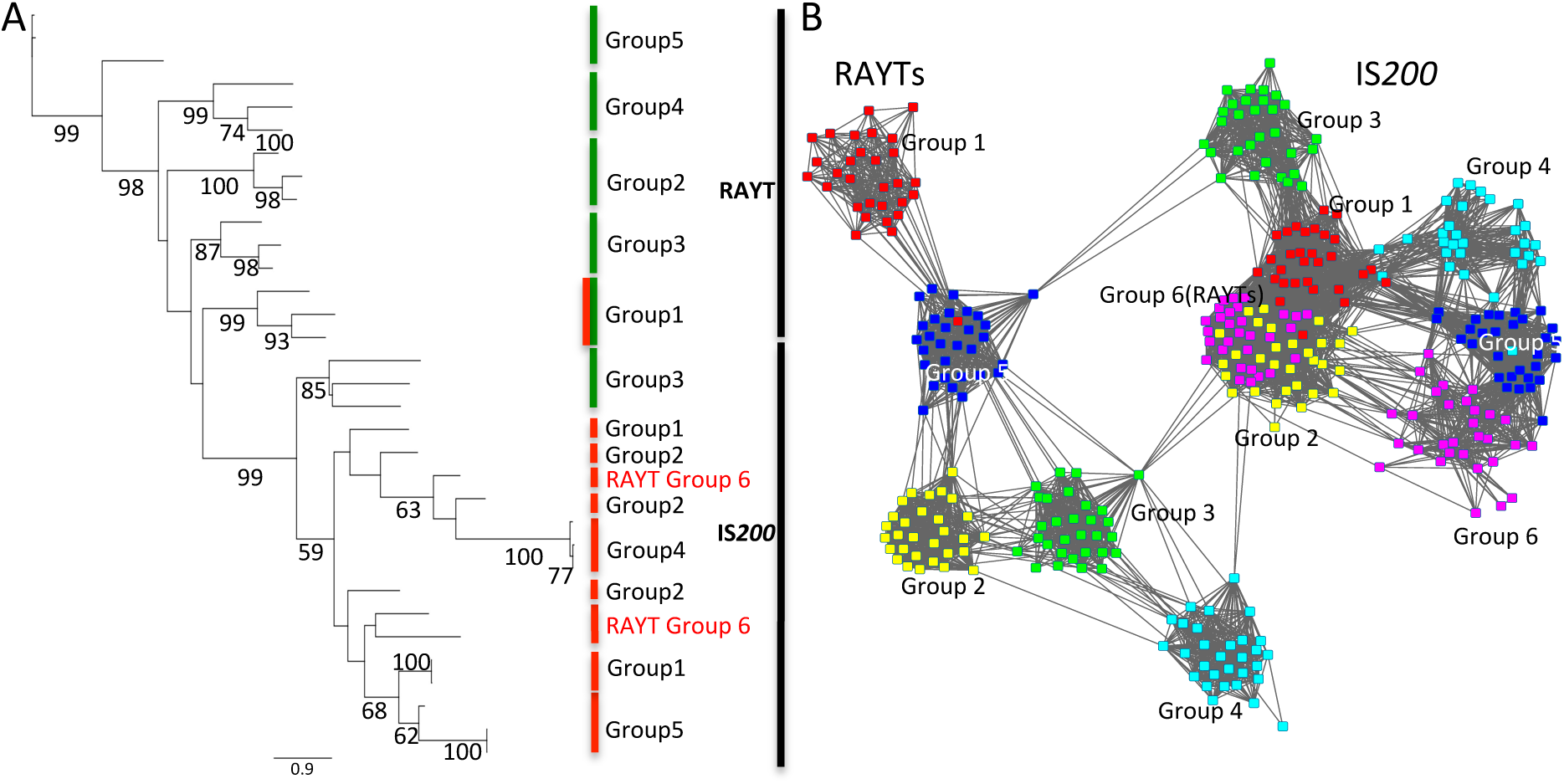
RAYTs and IS*200* gene families form monophyletic groups. (A) Phylogenetic tree. From each IS*200* and RAYT group, three random proteins were selected and aligned with MUSCLE (Edgar 2004). We excluded IS*200* Group 6, because the alignments could not be calculated correctly when members of this subgroup were included. The phylogeny was built using PHYML (Guindon and Gascuel 2003). All RAYTs are found in the same clade except for RAYT Group 6, which clusters within the IS*200* family. This is plausible considering the IS-like genome distribution characteristics. Green bars indicate low duplication rates (less than 5%); red bars indicate high duplication rate and high presence on plasmids (more than 17%). RAYT Group 1 is marked with a red/green bar as within the RAYT family it has the highest duplication rate (6.2%). (B) Protein similarity network of the RAYT (left) and IS*200* (right) gene family. We chose 30 random proteins from the six largest RAYT and IS200 groups and displayed them with Cytoscape as in Figures 3F and **4F**. RAYT Group 6 clusters within IS*200* Group 1 in line with the phylogeny in (A) and the genomic distribution characteristics in Figure 3.

The separation between RAYTs and IS*200* in the phylogenetic tree is not as clear in the protein network (Figure 5B). Although RAYTs seem to form relatively coherent clusters, these clusters do not include the RAYT-like Group 3 from the IS*200* family.

### Vertical transmission of RAYTs is supported by phylogenetic congruence at the sub genus taxonomic level

Given division of RAYT families into distinct groups it becomes possible to explore mode of transmission beyond simple inclusion or not on mobile elements such as plasmids as performed above. At the sub-genus level the phylogenetic trees of RAYT sequence groups 2 and 3 are very similar to the tree reconstructed from the two concatenated housekeeping genes (Figure 6) in contrast to the IS*110* phylogeny (**Supplementary Figure 8**). Group 2 agrees very well with the housekeeping phylogeny except for a few species. These incongruences could be caused either due to uncertainties at the level of the species tree (low bootstrap support) or rare horizontal gene transfer events. The phylogeny of Group 3 RAYTs is less clear, which is caused by frequent duplication and loss events. Nevertheless, sub-trees are consistent with the housekeeping phylogeny and RAYTs cluster within species. There is indication of one instance of horizontal transfer of a RAYT from *P. fluorescens* to *P. syringae* but the possibility that this is a rare duplication event that occurred in the ancestor of *P. fluorescens* and *P. syringae*, cannot be excluded.

**Figure 6.**
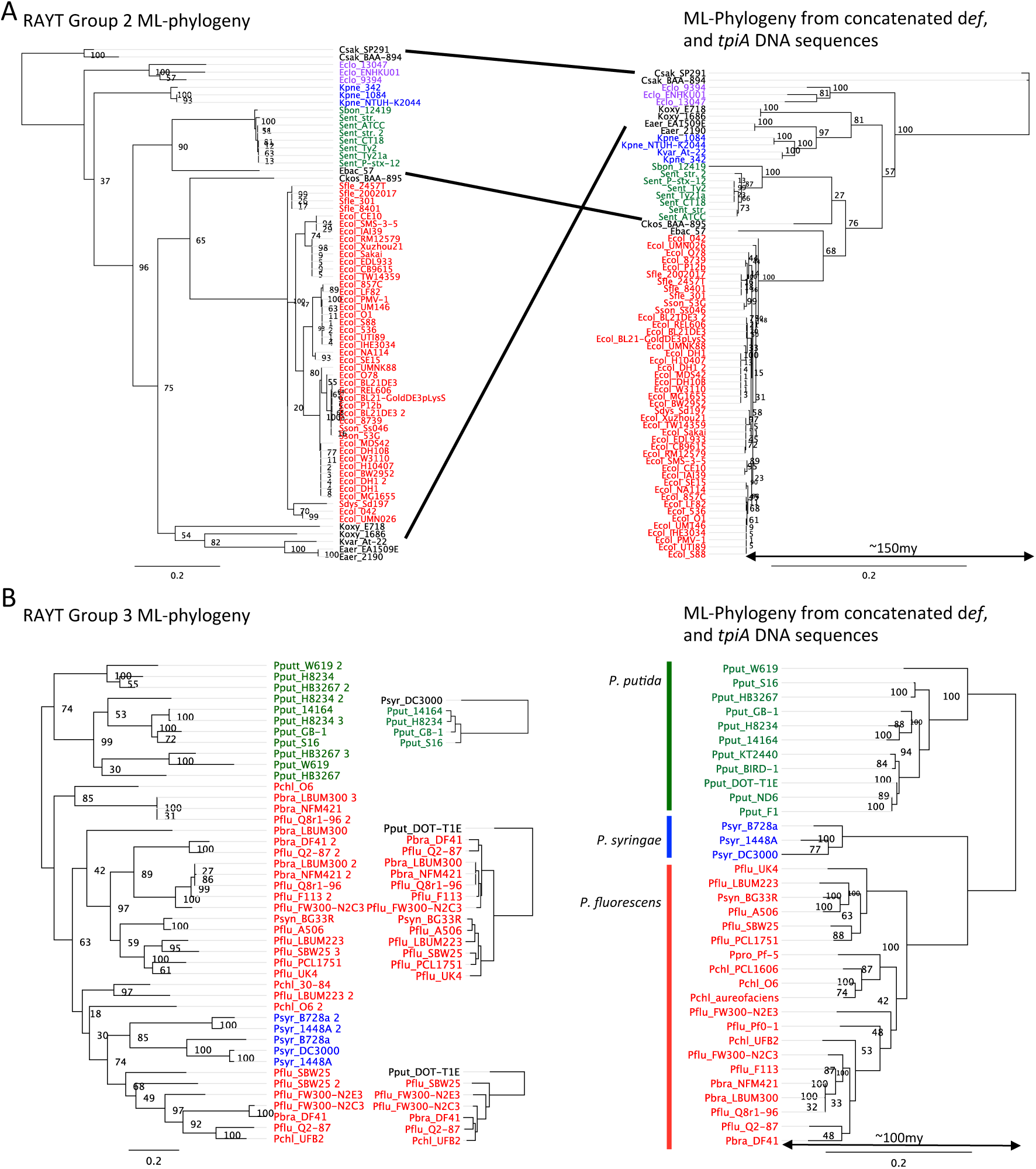
Comparison between RAYT phylogenies and housekeeping gene phylogenies shows vertical RAYT inheritance. (A) Shows the RAYT Group 2 phylogeny and its corresponding housekeeping gene tree of all RAYT Group 2 members that are found in the Enterobacteriaceae. *Escherichia, Salmonella, Klebsiella* and *Enterobacter cloacae* are coloured in red, green, blue and purple respectively. The remaining genera are in black and their corresponding positions in the housekeeping gene tree are connected with black lines. The most recent common ancestor is estimated to have lived about 150mya from a 16S rDNA tree **(Ochman & Wilson 1987)**. RAYT Group 2 genes occur mostly as single copy genes. (B) Shows the RAYT Group 3 phylogeny and its corresponding housekeeping gene tree for all RAYT Group 3 members that are found in *Pseudomonas* fluorescens, *P. putida and P. syringae.* They are coloured in red, green, and blue respectively. The most recent common ancestor is estimated to have lived about 100mya from a 16S rDNA tree **(Ochman & Wilson 1987)**. Small trees in between the housekeeping gene tree and the RAYT tree are corresponding housekeeping trees for the RAYT sub-trees. The strain name in black indicates the outgroup. RAYT Group 3 genes occur often in multiple copies per genome. Their evolution is dominated by gene loss and duplication events.

The phylogenies in Group 1, 4 and 5 contain fewer RAYTs. For Group 4 and 5 this is probably due to the much smaller family sizes. It is possible that the large diversity and low numbers per genus in Group 1 RAYTs is partly due to higher rates of horizontal gene transfer, which would also be supported by the presence of Group 1 RAYTs on plasmids. Group 4 and 5 RAYTs are less frequent but in the instances observed they evolve very slowly (low substitution rates compared to housekeeping genes) and their evolutionary histories are congruent with the housekeeping gene trees (**Supplementary Figure 8**).

### Over longer time scales horizontal gene transfer may dominate RAYT evolution

Over very long time scales (>100 million years) RAYT phylogenetic trees are not significantly more similar than IS phylogenies are to housekeeping gene trees (**Supplementary Table 1**). It is possible that this is due to horizontal gene transfer that becomes more likely over very long periods of time. It could also indicate that RAYTs first emerged from their IS*200*-like ancestors about 100-150 mya, possibly in *Pseudomonas* and *Enterobacteria* as these are the genera in which RAYTs are most commonly found. From these bacteria RAYTs could have been transferred very rarely to more distantly related bacteria.

## Conclusion

RAYTs are a recently described class of bacterial transposases that share a distant evolutionary relationship with the IS*200* gene family (Lam & Roth 1983; 1986; Barabas et al. 2008). The suggestion of a distant evolutionary relationship comes from the fact that both RAYTs and IS*200* share an identical catalytic centre (Bertels & Rainey 2011b; Nunvar et al. 2010; Ton-Hoang et al. 2012). Despite this similarity, the data presented here show that the RAYT and IS*200* families display different characteristics. Members of the IS*200* gene family show characteristics typical of ISs, whereas members of the RAYT family are, in many respects, almost indistinguishable from housekeeping genes. The housekeeping-like nature of RAYTs provides a firm basis for the earlier suggestion that RAYTs are domesticated transposons (Ton-Hoang et al. 2012; Siguier et al. 2014; Bertels & Rainey 2011a; 2011b).

The housekeeping role of RAYTs is unclear, but the existence of five distinct subfamilies (Figures 3A & 5B) raises the possibility of five different functions. Biochemical studies on RAYTs from *E. coli* provide evidence of *in vitro* endonuclease and DNA binding activity (Ton-Hoang et al. 2012). Such activity is thought to be integral to the capacity of RAYTs to catalyse transposition of non-autonomous REP / REPIN elements *via* an IS*200*-like mechanism (Barabas et al. 2008; Ton-Hoang et al. 2010). But this role alone is insufficient to explain the maintenance of RAYTs within genomes. In the absence of RAYTs catalysing their own transposition (for which there is no evidence), there is no mechanism for RAYTs to escape mutational decay, leading to the prediction that RAYTs must perform some function that benefits the host cell (Bertels & Rainey 2011a; 2011b).

Although there are various plausible scenarios, it is challenging to conceive of a single RAYT function that defines both its interaction with REPINs and its beneficial effect on the host bacterium. One possibility is that the various previously described (and likely co-opted) functions of REPs and REPINs, including effects on translation, transcription and DNA organization (Newbury et al. 1987; Yang & Ames 1988; Espéli et al. 2001; Voineagu et al. 2008; Liang et al. 2015), depend on some functional interaction with RAYTs. For example, binding of the RAYT protein to a particular REP or REPIN sequence may modulate its effects, with the different classes of RAYT possibly reflecting different specialisations in different bacterial groups. An alternate idea that we find intriguing — and which draws upon the endonuclease function of RAYTs — is that RAYTs function in concert with REPINs in a CRISPR-like way (Horvath & Barrangou 2010), with the palindromic REPIN structure acting as bait for incoming foreign DNA (Tobes & Pareja 2006; Treangen et al. 2009) that is then inactivated or degraded *via* the endonuclease activity of the RAYT. It is conceivable that such a scenario might also generate antagonistic co-evolution between bait and RAYT promoting the evolution of divergent RAYT subfamilies. Consistent with this idea is data from (Sinha et al. 2009) who reported that expression of the *E. coli* RAYT gene *yafM* is significantly elevated in cells in which competence is naturally induced. It is possible that over-expression prepares cell defences for the onslaught of foreign DNA.

**Supplementary Figure Legends**

**Supplementary Figure 1. Family size of ISs, RAYTs and housekeeping genes for four decreasingly stringent definitions of sequence relatedness.**

**Supplementary Figure 2. Same as Figure 2 except that all plots show data for four decreasingly stringent definitions of sequence relatedness.**

**Supplementary Figure 3. Same as Figure 3 except that all plots show data for four decreasingly stringent definitions of sequence relatedness.**

**Supplementary Figure 4. Same as Figure 4 except that all plots show data for four decreasingly stringent definitions of sequence relatedness.**

**Supplementary Figure 5. Protein alignment of a selection of 35 Group 2 RAYTs.** Boxed in red are conserved (>50%) and in blue not conserved putative DNA binding sites identified by (Messing et al. 2012).

**Supplementary Figure 6**. Protein alignment of all RAYT and IS*200* groups. Boxed in red are conserved (>50%) and in blue not conserved putative DNA binding sites identified by (Messing et al. 2012). Boxed in green are sites that are only conserved in the five RAYT groups and not in the IS*200* family.

**Supplementary Figure 7. Showing the different taxonomic distributions for all five RAYT groups as well as for each RAYT subgroup individually.** The radius of each pie slice indicates the proportion of sequenced species within the taxonomic class that contain a RAYT gene. The proportion of the pie indicates the proportion of species that contain a RAYT gene as a fraction of all RAYT genes in that group.

**Supplementary Figure 8. Sub-genus phylogenetic congruency analysis for RAYT Groups 1, 4, 5 and IS*110***.

**Supplementary Table 1. Phylogenetic congruency analysis of RAYT and IS containing species among the Gammaproteobacteria.**

**Supplementary Data. Nucleotide sequences for all RAYT groups.**

## ACKNOWLEDGEMENTS

We thank two anonymous referees for their critical review and helpful suggestions. FB acknowledges funding for a Ph.D. scholarship from the Allan Wilson Centre.

